# Deep Semi-Supervised Learning Improves Universal Peptide Identification of Shotgun Proteomics Data

**DOI:** 10.1101/2020.11.12.380881

**Authors:** John T. Halloran, Gregor Urban, David Rocke, Pierre Baldi

## Abstract

Semi-supervised machine learning post-processors critically improve peptide identification of shot-gun proteomics data. Such post-processors accept the peptide-spectrum matches (PSMs) and feature vectors resulting from a database search, train a machine learning classifier, and recalibrate PSMs using the trained parameters, often yielding significantly more identified peptides across *q*-value thresholds. However, current state-of-the-art post-processors rely on shallow machine learning methods, such as support vector machines. In contrast, the powerful training capabilities of deep learning models have displayed superior performance to shallow models in an ever-growing number of other fields. In this work, we show that deep models significantly improve the recalibration of PSMs compared to the most accurate and widely-used post-processors, such as Percolator and PeptideProphet. Furthermore, we show that deep learning is able to adaptively analyze complex datasets and features for more accurate universal post-processing, leading to both improved Prosit analysis and markedly better recalibration of recently developed database-search functions.

## Introduction

The field of proteomics has undergone explosive growth in the past two decades, largely fueled by technological breakthroughs in mass spectrometry. Most often accomplished through liquid-chromotography tandem mass spectrometry (LC-MS/MS) followed by a peptide-database search, proteomics experiments have concurrently seen rapid increases in the size and complexity of generated datasets, resulting in the proteome-scale analysis of whole biological systems^1,28,48^. In practice, however, the peptide-spectrum matches (PSMs) resulting from a database search are often *uncalibrated*, thus making qualitative comparisons difficult and diminishing overall identification accuracy. To overcome this, machine learning post-processors are widely used to *recalibrate* PSMs, greatly improving the yield of identifications at low false discovery rates (measured using *q*-values).

The most accurate of these post-processors are semi-supervised^6,24^, using decoy PSMs as negative training cases and examples gathered from target PSMs as positive training cases. For instance, one of the most popular post-processors, PeptideProphet,^6,26^ uses linear discriminant analysis (LDA) for fixed feature sets and pre-computed weights to recalibrate input PSMs. Another extremely popular post-processor, Percolator,^24^ adaptively learns feature weights by iteratively training a support vector machine (SVM) and recalibrating input PSMs using the final learned SVM parameters. Subsequent works have used shallow neural networks (i.e., Q-ranker^44^) and gradient boosted decision trees (i.e., Scavager^23^). However, these approaches limit analysis to specific search engines and sets of MS/MS features, similar to PeptideProphet. In contrast, *universal* post-processing–where arbitrary sets of MS/MS features may be analyzed accurately–has been made possible through Percolator’s adaptive algorithm.

By analyzing arbitrary feature sets and adaptively learning all parameters, universal post-processing has enabled two important developments for MS/MS analysis. The first is the recent development of machine learning methods which extract large numbers of sophisticated features from MS/MS datasets^13,16,17,46^ and rely on universal post-processing to use these complex feature sets for better PSM recalibration. The most notable of these feature extraction methods is Prosit,^13^ which uses deep neural networks to extract 64 informative features (per PSM) and feeds these features into Percolator to improve search results. The second is the quick and easy use of PSM recalibration by newly developed search algorithms. For instance, while established database-search scoring algorithms slowly adopted Percolator after its introduction (e.g., X!Tandem^49^ and Mascot^5^), recent search algorithms have rapidly adopted Percolator analysis and included it in initial software releases (e.g., XCorr *p*-values,^20^ DRIP,^15^ and combined res-ev *p*-values^32^).

However, while Percolator has demonstrated state-of-the-art performance compared to other popular post-processors,^47^ the use of shallow machine learning models (such as SVMs and gradient boosted decision trees) is potentially suboptimal given the cutting-edge performance exhibited by deep models^3^ across a large number of other fields. For example, deep learning models have greatly outperformed shallow methods and paved the way for recent groundbreaking advances in computer vision,^30,34^ natural language processing,^10,39^ genomics,^2,50^ particle physics,^4^ climate analysis,^45^ speech recognition,^18,19^ and medical diagnosis.^33,36,43^

In this study, we explore whether using the powerful learning capabilities of deep neural networks (DNNs) further improves PSM recalibration and MS/MS post-processing accuracy. Compared to the shallow models used in the state-of-the-art universal post-processor Percolator, we demonstrate that the use of DNNs allows more peptide identifications (for all considered *q*-value thresholds) across a large number of diverse datasets: nine high-res MS1/MS2 datasets collected using six different protein databases. These benchmarks include Prosit and Comet feature sets, as well as PSM features collected from a recently developed scoring algorithm (`NegLog10CombinePValue`) designed for new MS/MS machines and datasets.^32^

Most importantly, we show that DNNs provide the largest performance gains for datasets which contain the most information for machine learning classification. For instance, the collected `NegLog10CombinePValue` features contain over 50% more PSM information (measured using *mutual information*^7^) than Comet features for the searched MS/MS spectra and databases. At a 1% *q*-value threshold, DNNs nearly double the average recalibration accuracy for these highly informative `NegLog10CombinePValue` features compared to using SVMs in Percolator (for one of these datasets, Percolator fails to improve recalibration altogether). This thus demonstrates that, compared to the increased learning capabilities of deep models, shallow methods are suboptimal at exploiting information-rich MS/MS datasets for improved peptide identification. Furthermore, we show that the use of DNNs drastically improves PSM recalibration compared to the other shallow models used in non-adaptive post-processors Scavager,^23^ Q-ranker,^44^ and PeptideProphet,^26^ leading to an average 14.2% increase in peptides identified at a strict *q*-value of 1% for data collected from the widely-supported Comet search engine.^12^

In what follows, we refer to the universal post-processor which employs DNNs for deep semi-supervised learning as *ProteoTorch DNN*.

## Results

### Deep Semi-Supervised Learning Improves Peptide Identification Accuracy

ProteoTorch DNN utilizes an iterative, semi-supervised training algorithm (illustrated in Figure 1). At each iteration, decoys and statistically significant targets (measured using *q*-values) are assigned negative and positive training labels, respectively. The parameters of a DNN with three hidden layers (each with 200 neurons) are learned using the training labels and accompanying PSM features, and are then used to score PSMs. Subsequently, these scores are used to calculate new target *q*-values and the training process repeats. Importantly, the training and testing sets used during the algorithm are kept disjoint using three-fold cross-validation,^14^ thus preventing the overfitting of learned parameters. Further details of this deep semi-supervised algorithm are described in Methods.

**Figure 1:**
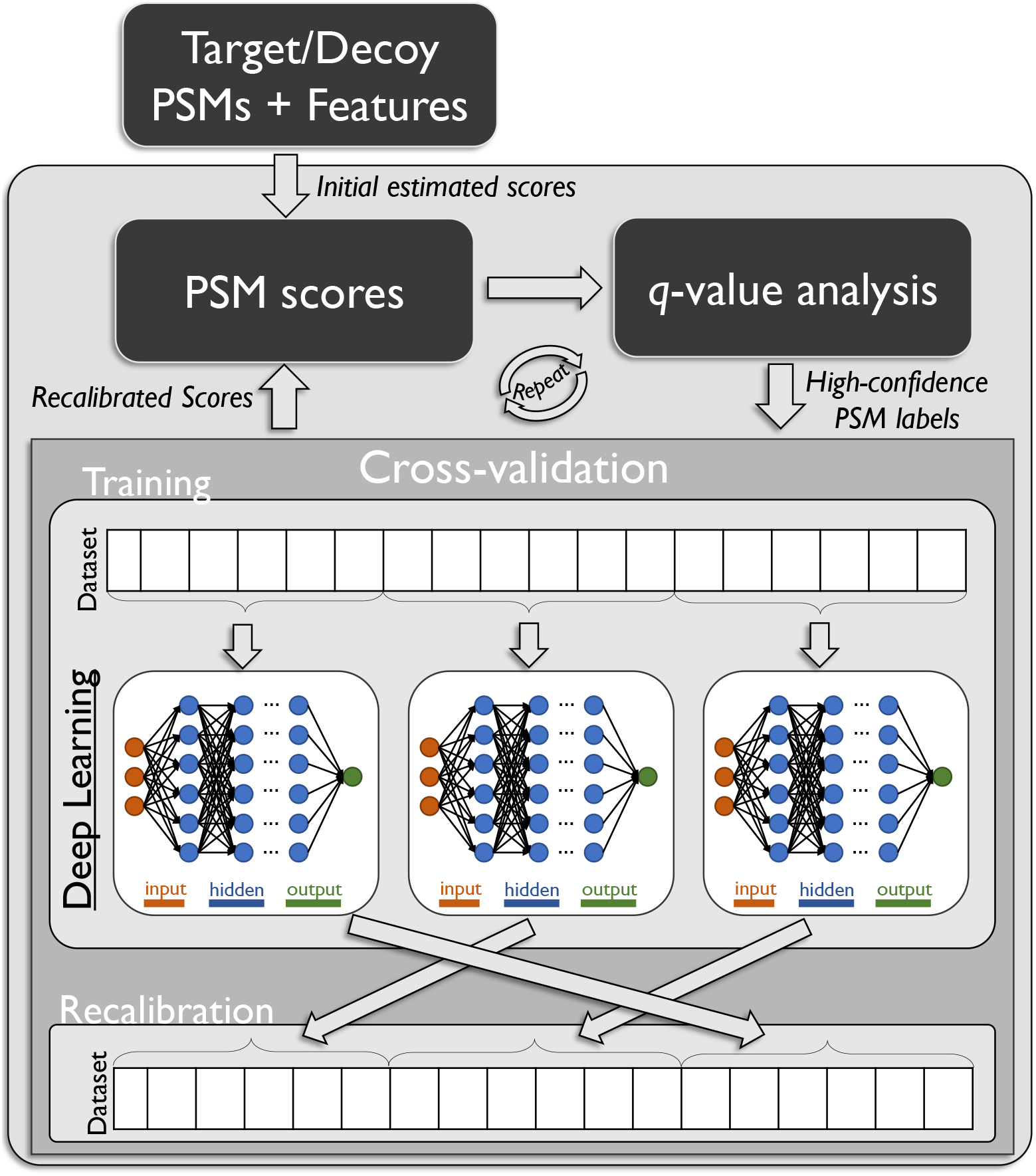
Semi-supervised deep learning algorithm. Target and decoy PSMs, with corresponding feature vectors, are input and used to estimate initial PSM scores. These scores are used to calculate *q*-values and determine highly-confident (given a *q*-value threshold) target PSMs for training. The dataset is split into separate cross-validation (CV) bins, and the PSMs in each bin (i.e., decoys and high-confidence targets) are used to train a deep neural network classifier. Each trained classifier is then used to rescore PSMs in a separate CV bin, and this process repeats either til convergence or a user-specified number of iterations has passed.

To assess universal post-processing performance using DNNs, nine benchmark datasets were evaluated: three Prosit datasets, three datasets collected using the recently developed `NegLoglO-CombinePValue` database-search function (which is the calibrated combination of highly accurate, *p*-value scoring functions), and three datasets collected using the widely-used Comet data-basesearch algorithm. Prosit datasets were derived searching the same human gut spectra with three increasingly-complex protein databases, and acquired from the online data repository of.^13^ `NegLog10CombinePValue` datasets were collected by searching three different collections of high-resolution MS/MS data: spectra collected from the gargle solutions of COVID-19 patients^22^ (referred to as COVID-19); spectra collected from human red blood cells infected with the malaria parasite *Plasmodium falciparum* (referred to as Plasmodium^42^); and 24,931,642 spectra from a draft of the human proteome (referred to as Human^28^). Comet datasets were collected by searching the same MS/MS spectra and protein databases as `NegLog10CombinePValue`. All MS/MS data details and Crux `NegLog10CombinePValue` and Comet search settings are described in Methods. The size of these datasets range from slightly over one-million to over twenty-million PSMs. The resulting number of PSMs, number of features, and utilized protein databases for each dataset are listed in Table 1.

**Table 1:** Benchmark datasets

| Data set | Species | Search engine | # of features | # of PSMs |
| --- | --- | --- | --- | --- |
| Prosit Human+Bacteria<sup>40</sup> | Human and Bacteria | Andromeda<sup>8</sup> | 64 | 4,264,003 |
| Prosit All<sup>40</sup> | All organisms | Andromeda | 64 | 4,325,290 |
| Prosit IGC+All<sup>40</sup> | IGC and all organisms | Andromeda | 64 | 4,643,570 |
| COVID-19<sup>22</sup> | Human and SARS-COV-2 | Crux tide-search | 21 | 1,336,153 |
| Plasmodium<sup>42</sup> | *P. falciparum* | Crux tide-search | 21 | 1,336,386 |
| Human<sup>28</sup> | Human | Crux tide-search | 21 | 19,961,754 |
| COVID-19<sup>22</sup> | Human and SARS-COV-2 | Comet | 21 | 1,363,163 |
| Plasmodium<sup>42</sup> | *P. falciparum* | Comet | 21 | 1,622,376 |
| Human<sup>28</sup> | Human | Comet | 21 | 20,321,131 |

Prosit and Crux `NegLog10CombinePValue` datasets are only supported for recalibration by universal post-processors ProteoTorch DNN and Percolator. Comet searches were supported, and recalibrated, using all evaluated post-processors, i.e., ProteoTorch DNN, Percolator, PeptideProphet, Q-ranker, and Scavager. Settings for all post-processing methods were chosen to ensure as fair comparison of recalibration performance as possible (see Methods). For all evaluated methods, *q*-values were calculated using target-decoy competition.^11^ The post-processing performance for all datasets is plotted in Figure 3, and the number of significant PSMs identified at *q*-value threshold 1% per dataset is listed in Table 2.

**Table 2:** Number of significant PSMs identified at a *q*-value threshold of 1% for the datasets listed in Table 1. For all methods, *q*-values were measured using target-decoy competition. Database-search functions `NegLog10CombinePValue` and XCorr are listed for Crux and Comet datasets, respectively. For post-processors Q-ranker, PeptideProphet, and Scavager, dashes denote the data’s feature set is not supported. The largest number of significant PSMs per dataset is highlighted in bold.

| Data set | ProteoTorch<br>DNN | Percolator | Q-ranker | PeptideProphet | Scavager | NegLog10CombinePValue | XCorr |
| --- | --- | --- | --- | --- | --- | --- | --- |
| Prosit Human+Bact. | **34,055** | 33,350 | – | – | – | – | – |
| Prosit All | **35,796** | 34,963 | – | – | – | – | – |
| Prosit IGC+All | **107,118** | 101,241 | – | – | – | – | – |
| Crux COVID-19 | **36,636** | 35,419 | – | – | – | 32,666 | – |
| Crux Plasm. | **106,036** | 99,491 | – | – | – | 92,833 | – |
| Crux Human | **1,584,028** | 1,527,775 | – | – | – | 1,532,936 | – |
| Comet COVID-19 | **32,489** | 31,410 | 29,658 | 23,200 | 28,460 | – | 15,812 |
| Comet Plasm. | **132,993** | 130,803 | 126,576 | 121,379 | 116,153 | – | 101,328 |
| Comet Human | **1,700,926** | 1,668,551 | 1,553,954 | 1,545,288 | 1,480,338 | – | 1,315,507 |

Across all other evaluated *q*-values, semi-supervised learning using DNNs achieves more significant identifications than using SVMs in Percolator for all post-processing tasks: Prosit analysis, recalibration of `NegLog10CombinePValue` search results, and recalibration of Comet search results. For the Prosit dataset generated using the largest protein database (IGC+All, collected by searching over ten-million protein sequences), the use of deep models increases the number of significant PSMs over SVMs in Percolator by 5.8% at a *q*-value of 1%. For the complex search function `NegLog10CombinePValue`–developed to accurately compute *p*-values for high-resolution MS/MS data–DNNs increase the number of significant identifications by 8.2% on average when using a strict *q*-value threshold of 1%, whereas the use of shallow models in Percolator only results in an average increase of 4.1%. For the post-processing of Comet search results, the use of DNNs out-performs all other methods for all evaluated *q*-values, identifying an average 14.2% more peptides at *q*-value tolerance 1% than the non-adaptive methods Scavager, Q-ranker, and PeptideProphet, which rely on shallow machine learning models.

### Deep models better exploit MS/MS information than shallow models

To study the amount of information available to machine learning classifiers during semi-supervised learning, the average *mutual information* between each PSM feature and target/decoy label was calculated for each MS/MS dataset. Mutual information is the information-theoretic quantity which measures the information one random variable provides about another; when two random variables are independent, their mutual information is zero. Otherwise, higher mutual information denotes higher dependence. Thus, higher values of average mutual information denote higher amounts of available information in a dataset detailing whether a PSM is a target or a decoy. Note that mutual information is always nonnegative.

In Figure 2(a), the average mutual information per feature and target/decoy label is compared between the three different Prosit datasets, where the same MS/MS spectra were searched with three different protein databases. In Figure 2(b), average mutual information is compared between the three pairs of Crux `NegLog10CombinePValue` and Comet datasets, where the same spectra and protein databases were searched using both Crux and Comet to produce different feature sets. For Prosit results, the dataset generated using the largest database, IGC+All, contains the most informative feature values, providing over 50% more information than each other dataset searched (i.e., a minimum 50.5% and maximum 60.7% more information than the other datasets collected using smaller protein databases). For Crux and Comet results, the Crux `NegLog10CombinePValue` feature sets provide an average 53% more information for classification than corresponding Comet feature sets generated searching each of the COVID-19, Plasmodium, and Human data.

**Figure 2:**
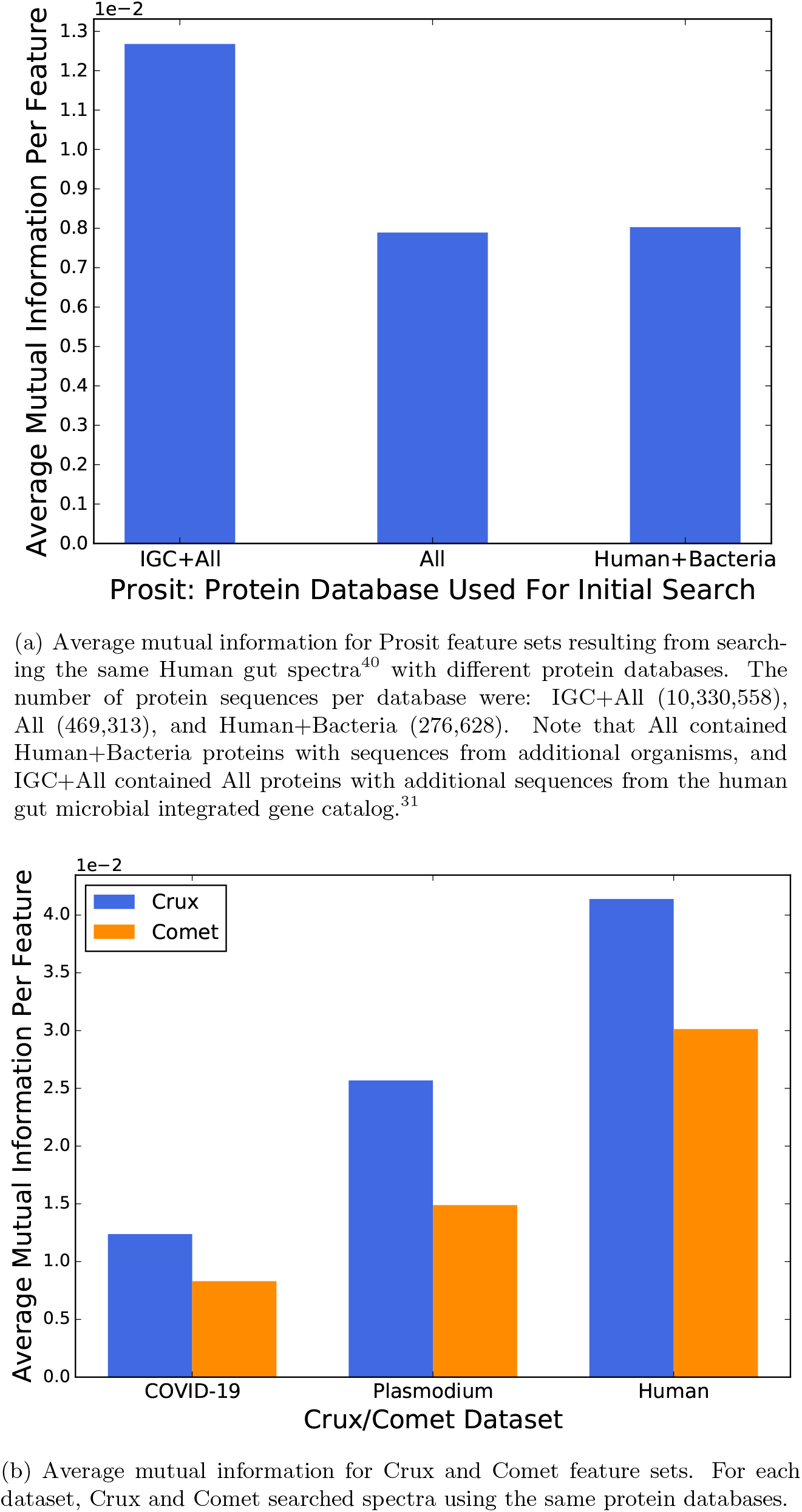
Average mutual information for each dataset in Table 1. The average mutual information displays the mean amount of information each feature provides for whether a PSM is a target or a decoy.

As seen in Figures 3(a)-3(c), while DNNs generally improve Prosit accuracy over Percolator, this improvement is most pronounced for the IGC+All dataset which contains substantially more information than the other two Prosit datasets. This trend is also true for the Crux and Comet results; Percolator effectively improves Comet post-processing, but the SVMs used are much less effective at exploiting the larger amount of information available in `NegLog10CombinePValue` features, failing to recalibrate PSMs for the Crux Human dataset. In contrast, the increased training capabilities of deep models effectively use larger amounts of information to better classify target and decoy PSMs, allowing ProteoTorch DNN to successfully recalibrate PSMs for all `NegLog10CombinePValue` datasets.

**Figure 3:**
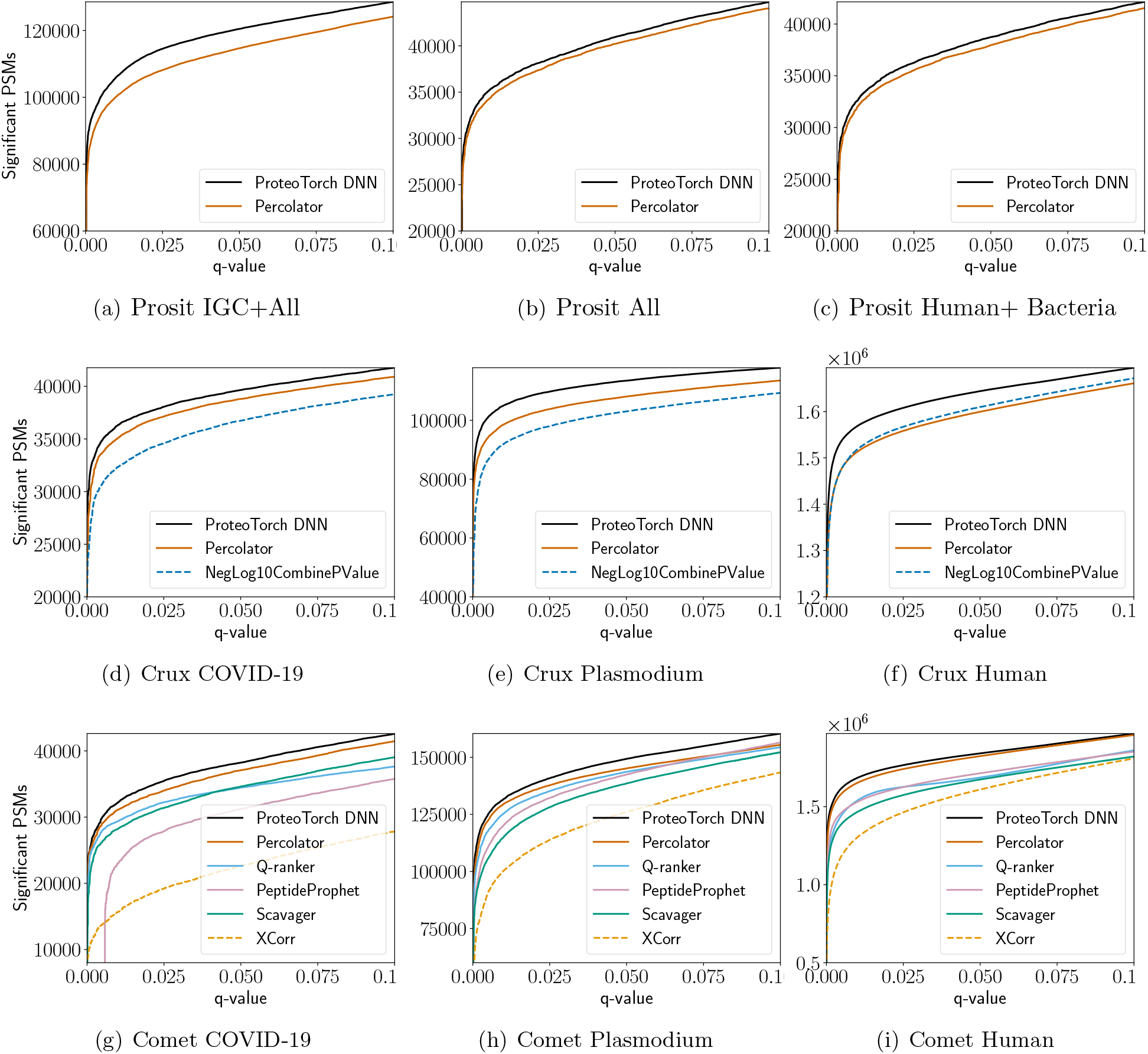
Recalibration performance plots for the Prosit, Crux tide-search, and Comet benchmark datasets listed in Table 1. The *x*-axis corresponds to *q*-values and the y-axis displays the number of PSMs deemed significant at each *q*-value. Target-decoy competition^11^ was used to compute *q*-values for all methods. Prosit and Crux `NegLog10CombinePValue` post-processing are currently only supported by universal post-processors ProteoTorch and Percolator. Underlying search function performance for Comet’s XCorr and Crux’s `NegLog10CombinePValue` are additionally plotted using dashes.

## Discussion

This was the first study to explore the impact of recent deep learning advances for semi-supervised post-processing of shotgun proteomics results. Across all datasets and evaluated *q*-values, the use of DNNs outperforms the shallow models used in the universal post-processor, Percolator. This increase in performance is markedly pronounced for complex datasets with more information available to machine learning classifiers during semi-supervised learning. For the `NegLog10CombinePValue` feature sets-which, compared to corresponding Comet feature sets, contain more classification information for each MS/MS dataset-the shallow networks used by Percolator have difficulty improving discrimination between calibrated target and decoy PSM scores (even failing to improve performance for the Crux Human dataset). In contrast, DNNs are capable of using these highly informative features to better distinguish target and decoy PSMs, resulting in nearly double the recalibration improvement, at a strict *q*-value threshold of 1%, compared to the use of SVMs in Percolator. Furthermore, for Prosit IGC+All-which contains significantly more information for MS/MS classification compared to the other Prosit datasets-deep models are able to better differentiate targets from decoys, identifying 5.8% more PSMs than Percolator at a *q*-value of 1%.

For Comet searches, post-processing performance using DNNs largely outperforms all other methods, identifying an average 14.2% more peptides at a 1% *q*-value than the non-adaptive methods Q-ranker, PeptideProphet, and Scavager. For the non-adaptive method PeptideProphet, it is also worth noting the effect of using precomputed LDA weights across different datasets. While PeptideProphet improves XCorr accuracy across all *q*-values for the datasets Comet Plasmodium and Comet Human, no post-processed matches are deemed significant at stringent (near zero) *q*-values for Comet COVID-19, which leads to worse performance than simple XCorr (i.e. without post-processing) for this low *q*-value range. This demonstrates that the use of precomputed weights prevents effective generalization across the different datasets that were evaluated, and even degrades Comet search accuracy over one of the datasets at very strict *q*-values.

We’ve thus shown that deep models may be used to significantly improve the recalibration of MS/MS database-search results compared to shallow models, which are used in the most accurate post-processing packages in widespread use. In particular, deep semi-supervised learning was demonstrated to effectively adapt to diverse MS/MS data and feature sets, thus enabling state-of-the-art universal post-processing across a variety of downstream tasks: for Prosit analysis, recalibration analysis of a high-resolution *p*-value search function, and recalibration of Comet search results. Furthermore, deep models provide markedly better recalibration performance than available shallow models for MS/MS datasets with high amounts of detailed PSM information.

Having established that deep models may be used to significantly improve MS/MS post-processing analysis, several important avenues of future work are available. One such avenue is studying the impact the demonstrated improvements in peptide identification have on protein identification accuracy, assessed using different statistical strategies such as the picked protein false discovery rate.^41^ Furthermore, it is clear that the amount of mutual information contained in MS/MS datasets plays an important role in machine learning classification accuracy and potential recalibration performance. Thus, exploration of what experimental factors (e.g., search engine, protein database statistics, searching settings, etc.) produce the most information-rich MS/MS datasets will help us understand when deep models may provide the greatest performance gains for universal post-processing. Finally, the exploration of different neural architectures for other widely used feature sets may lead to further improvements in peptide identification. We note that this last line of work may be efficiently performed using the ProteoTorch DNN software, which is accessible across all major platforms (Windows, Linux, and Mac) and freely available at github.com/proteoTorch/proteoTorch.

## Methods

### Deep learning model and semi-supervised training process

ProteoTorch uses an iterative, semi-supervised training procedure to recalibrate input targets and decoy PSMs (illustrated in Figure 1). By construction, all decoy PSMs are incorrect identifications, and are thus assigned negative labels, whereas positive training examples are estimated as the set of target PSMs with scores achieving a stringent, user-specified *q*-value. These scores are then re-evaluated in each training iteration and the positive label assignments are updated, using the predictions of a classifier that is trained to distinguish between positively and negatively labeled PSMs. This overall process repeats either for a user-specified number of iterations or until convergence. Furthermore, to prevent overfitting and to improve generalizability, three-fold cross-validation (CV) is carried out in the overall procedure, where the dataset is partitioned into three separate test and train splits (as described in^14^). Thus, PSMs from a training set are always disjoint from the corresponding test set (i.e., the set of PSMs to be re-scored). When the classifier is a linear SVM,^25^ the described training scheme is equivalent to Percolator.^24^

The DNN classifier used in ProteoTorch is a Multilayer Perceptron, which is the most suitable deep learning architecture for this problem as the input PSM features have no regular (or temporal) structure that could be exploited by convolutional or recurrent neural networks. To avoid confusion in the DNN training details that follow, we distinguish the use of the term *iteration* to refer to a single step in the overall semi-supervised training procedure, and the use of the term *epoch* to refer to a single pass through the data when training the DNN–thus, each iteration consists of several epochs of training within a CV fold, and in turn, each epoch consists of many gradient update steps of small batches of training data.

By default, the DNN consists of three hidden layers with 200 ReLU (rectified linear unit) neurons and is trained for 50 epochs using the Adam^29^ optimization algorithm. Within an iteration, DNN training starts with a learning rate of 0.001, which is periodically reduced (ten times) by a factor of up to 50 over the course of five training epochs and reset again. At the low-point of each of these ten reduction cycles, a snapshot model^21^ is taken. The resulting ten snapshot models are combined into one final ensemble model, while taking into account the validation accuracy of the individual models, and this ensemble model serves as the trained classifier for the iteration. This snapshot-based process has the distinct advantage of producing a reasonably diverse ensemble classifier without requiring additional training time. The loss function used during training is a modified cross entropy loss, which increases the loss incurred by false positive predictions by a factor of four (leading to higher penalties for incorrectly predicted training decoys). Additionally, label smoothing is used, which effectively shifts the target probability for each class slightly away from 100%. These loss function modifications were implemented to account for the asymmetry in the reliability of the labels in PSM datasets.

Prior to the first iteration, a large ensemble of snapshot models (30 by default) is used to estimate initial PSM scores, which decreases the number of iterations necessary for overall convergence. Furthermore, to regularize training with coarse initial scores, the dropout rate in the first iteration is set to 0.5. As the estimated scores improve in subsequent iterations, the benefit of dropout lowers significantly and, thus, the dropout rate is set to zero for all iterations beyond the first.

All discussed DNN hyperparameters are the default values used for experiments.

### Datasets and Processing

Database search results were collected using Crux^35^ and Comet^12^ for three different high-resolution MS1/MS2 datasets: a SARS-COV-2 dataset collected from COVID-19 patients,^22^ downloaded from the PRoteomics IDEntifications (PRIDE) database (project PXD018682); a *Plasmodium falciparum* dataset,^42^ downloaded from MassIVE [MSV000084000](https://doi.org/10.25345/C5BD2H); and a draft of the human proteome,^28^ downloaded from PRIDE project PXD000561. COVID-19 and Plasmodium datasets were searched with Crux using the mzML spectra files supplied in the respective data repositories, and converted to ms2 format using msconvert^27^ for Comet searches. RAW human files were converted to ms2 format using msconvert with peak-picking and deisotoping filters. Plasmodium falciparum proteins were downloaded from PlasmoDB on April 26, 2020, resulting in 14,722 sequences. For the COVID-19 dataset, both human and SARS-COV-2 proteins were searched by combining the UniProt Human reference proteome and SARS2 organism proteins (https://covid-l9.uniprot.org), both accessed April 30, 2020 and resulting in 20,363 sequences. The database used to search the human dataset was the same Uniprot human reference proteome used for the COVID-19 data.

All searches were fully tryptic, allowed two missed cleavages, specified a fixed modification of carbamidomethylation cysteine, and concatenated target-decoy results. The COVID-19 and human datasets were searched using a precursor tolerance of 10 ppm and fragment mass tolerance of 0.05 Da, while the Plasmodium dataset was searched using a precursor tolerance of 50 ppm and fragment mass tolerance of 0.02 Da. For searches of the COVID-19 dataset, methionine oxidation and aspartic acid deamidation were specified as variable modifications. For searches of the Plasmodium dataset, a variable peptide N-terminal pyro-glu modification was specified and, for Comet searches, a variable protein N-terminal acetylation was further specified (this variable modification was not yet supported in the utilized version of Crux). For the large-scale human dataset (consisting of 426 files and 24,931,642 total spectra), a variable methionine oxidation modification and a variable N-terminal glutamine cyclization were specified, as well as a variable protein N-terminal acetylation for Comet searches.

Crux `tide-search` was run with `--exact-p-value true --score-function both`, thus ranking PSMs by `NegLog10CombinePValue` (i.e., the calibrated combination of XCorr^20^ and res-ev^32^ *p*-values) during database search. Comet’s `num_output_lines` parameter was set to five (matching Crux `tide-search’s` default). Both Comet and Crux searches were set to output pepXML and PIN (Percolator INput) files. All Crux and Comet settings not discussed were left to their default values. For each dataset, resulting pepXML files were combined using `InteractParser` from the Trans-Proteomic Pipeline^9^ (resulting PIN files were similarly concatenated using Linux command-line utilities). Per dataset, the combined pepXML files were subsequently converted to tsv format (for Q-ranker processing) using Crux `psm-convert`.

Prosit PIN files were downloaded from PRIDE project PXD010871. As described in,^13^ these three PIN files are the result of Prosit post-processing Andromeda searches over the same human gut MS data^40^ with three increasingly complex protein databases: (1) Human+Bacteria (276,628 proteins), (2) All organisms (469,313 proteins), and (3) IGC+All, which included the human gut microbial integrated gene catalog^31^ (IGC, 10,330,558 proteins).

The described benchmark files are summarized in Table 1.

### Software details

ProteoTorch is implemented in Python, uses `PyTorch`^37^ as its deep learning backend, and is available for download at https://github.com/proteoTorch/proteoTorch. Crux version 3.2-2372dae was used for combined res-ev *p*-value searches using `tide-search`, Q-ranker post-processing using q-ranker, and conversion of pepXML files to tsv format using psm-convert. Additionally, a patch was necessary to prevent Q-ranker from crashing while writing recalibrated scores, available at https://github.com/johnhalloran321/crux-toolkit. Comet version 2019.01 rev. 5 was used for all Comet searches. Percolator post-processing was performed using version 3.05. Peptide-Prophet post-processing was run using the Trans-Proteomic Pipeline^9^ version v5.2.0 Flammagenitus. Scavager post-processing was performed using version 0.2.1a3.

Unless specified otherwise, all post-processing parameters were left to their defaults. Percolator and ProteoTorch were run using target-decoy competition^11^ with flags `-Y` and `--tdc true`, respectively. Additionally, Percolator was run with flags `-m $targetfile -M --decoyfile -U`, thus reporting both target and decoy results and skipping taking the maximum-scoring PSM per peptide. PeptideProphet was run in the semi-supervised mode and reporting of all recalibrated target/decoy PSM scores, i.e., with flags `ZERO DECOY=DECOY_ DECOYPROBS`. To ensure methods utilized the same set of input features for as fair of a comparison of recalibration performance as possible, retention time prediction was not used in post-processors supporting this option (Percolator and Scavager).

The average mutual information per MS/MS dataset was calculated using `mutual_info_classif` from `scikit-learn`^38^ version 0.22.2.

### Hardware details

All described database searches and post-processing results were run on the same machine with an Intel Xeon E5-2620, 64GB RAM, and an NVIDIA Tesla K40 GPU.

## Data Availability

Database search and post-processing results used to perform all analysis and generate all figures are available at http://jthalloran.ucdavis.edu/proteoTorchData.html.

## Code availability

ProteoTorch DNN is available for download at github.com/proteoTorch/proteoTorch. *q*-value analysis plots were created using the ProteoTorch module proteoTorchPlot, with the number of PSMs identified at specific *q*-values also output by this module. The code used to calculate mutual information is packaged with the data.

## Acknowledgments and Funding

The work of JH and DR is in part supported by the National Center for Advancing Translational Sciences (NCATS), National Institutes of Health, through grant UL1 TR001860 and a GPU donation from the NVIDIA Corporation. The work of GU and PB is in part supported by grants NSF NRT 1633631 and NIH GM123558 to PB.

## Author Contributions

JH and PB conceived of the project. GU designed the neural architecture and wrote all deep learning code. JH wrote the general semi-supervised learning package, with substantial input from GU. JH acquired the data, performed the analyses, and tested hyperparameters. DR acquired the compute resources for the presented experiments. JH and GU wrote the paper, with edits from PB.

